# Site-specific O-glycans influence lacritin structure and multimerization in tears

**DOI:** 10.64898/2026.03.30.715376

**Authors:** Vincent Chang, Ryan J. Chen, Isaac Lian, Keira E. Mahoney, Jeff Romano, Gordon Laurie, Stacy A. Malaker

## Abstract

Lacritin is an abundantly expressed glycoprotein in tear fluid and plays key roles in immune response, tear secretion, and bacterial killing. These biological functions are tightly regulated through several biochemical mechanisms including multimerization, proteolysis, and alternative splicing, especially within its C-terminal domain. Given its critical role at the ocular surface, lacritin is currently under investigation as a diagnostic biomarker and therapeutic candidate for dry eye disease (DED). However, despite over three decades since its initial discovery, the functional significance of the O-glycans that comprise more than 50% of its molecular weight remain largely unknown. To address this gap, we leveraged mass spectrometry (MS)-based glycoproteomics and molecular dynamics (MD) to explore the structural role of site-specific O-glycans on C-terminal lacritin. In doing do, we identified distinct glycosylation profiles between monomeric and multimeric lacritin, particularly at glycosites located near crosslinking residues (Lys101 and Lys104) that modulate multimer formation. Building on our glycoproteomics data, we performed MD simulations on monomer and multimer glycoforms and revealed that O-glycans participate in intra-glycan-protein interactions, thereby affecting the conformational flexibility of lacritin and the spatial arrangement of Lys101 and Lys104. Finally, we quantified the solvent-accessible surface area (SASA) of Lys101 and Lys104, highlighting that proximal O-glycosylation is predicted to affect the propensity of these residues to participate in crosslinking. Taken together, these findings underscore a central role for lacritin O-glycans in affecting structural topology with implications for its downstream biological activity.

## Introduction

Lacritin is a ∼23 kDa glycoprotein at the ocular surface that participates in tear production^1–3^, antimicrobial activity^4,5^, epithelial regeneration^6,7^, and protection against inflammatory stress^8^. Although lacritin is highly abundant (∼18 to 27 μM)^9^ in healthy tear fluid, its expression levels are known to be dysregulated in ocular pathologies such as dry eye disease (DED), an affliction experienced by over 400 million individuals worldwide.^9^ Given its ubiquitous role in tear film, lacritin has been studied extensively in the past 30 years and is currently under investigation as a diagnostic biomarker and therapeutic candidate for DED.^10^

Recent advances in lacritin-based diagnostics and therapeutics are largely driven by foundational biochemical discoveries that elucidated the mechanistic basis of lacritin function. For instance, Ma et al. first uncovered that the downstream biological activity of lacritin is mediated through its binding interaction with syndecan-1 (SDC1), which initiates calcium signaling involving a G-protein coupled receptor (GPCR).^11^ Importantly, signaling through the lacritin-syndecan-1 axis is established through noncovalent interactions between the C-terminal alpha helix of lacritin (residues 114 to 138) and the conserved GAGAL N-terminal domain of SDC1.^12^ Additionally, 3-O sulfation (generated following heparanase-mediated deglycanation of SDC1) and chondroitin sulfate within the N-terminal region of SDC1 are required for lacritin binding.^11,13^ These findings have since informed the development of Lacripep™, a peptide-based therapeutic candidate for DED derived from the lacritin C-terminus which aims to reverse the pathological outcomes of dry eye.^10,14^

Several regulatory mechanisms influence the ability of lacritin to bind SDC1. For instance, lacritin forms dimers, trimers, and multimers through transglutaminase (TGM2)-mediated crosslinking, where Lys101 and Lys104 act as donor residues and Gln125 acts as an acceptor residue.^15^ Although the biological role of multimers in circulating tear fluid is still unknown, the process of multimerization significantly impairs lacritin binding to SDC1 and thus abrogates downstream signaling. Alternative splicing to generate lacritin isoforms A, B, C, and D is also known, where A represents the canonical sequence and is the only isoform known to bind SDC1.^2^ Until recently, isoform D was not predicted to exist at the protein level, and it still remains to be seen whether isoform D can participate in multimerization.^16^ Finally, several studies have demonstrated that C-terminal proteolysis of lacritin generates cleavage-potentiated fragments which can engage SDC1 or participate in bacterial killing.^5,14,17,18^ Collectively, these processes contribute to ocular homeostasis through regulating lacritin effector function.

Crucially, lacritin is also highly decorated with glycans that contribute over 50% of its molecular weight.^16,19^ Glycosylation is a post-translational modification (PTM) whereby glycans most commonly covalently modify Asn (N-linked) and Ser/Thr residues (O-linked) in a non-templated fashion.^20–22^ These glycan modifications can alter the physiochemical properties of the underlying protein to influence protein folding, confer proteolytic resistance, and shape protein-protein interactions.^21–25^ However, despite the biochemical significance of glycosylation, its functional role in lacritin biology remains largely unexplored. This gap in understanding is primarily attributable to experimental difficulties in studying lacritin glycans. Most notably, low expression yields in mammalian systems have prevented analysis by controlled biochemical assays, while the analytical complexity of tear fluid has prevented its characterization by MS-based glycoproteomics.^26,27^

Moreover, the substantial heterogeneity inherent to extensively glycosylated proteins and the high conformaitonal flexibility of glycans has long hindered structural elucidation efforts by X-ray crystallography or cryogenic electron microscopy (cryo-EM).^28^ Although molecular dynamic (MD) simulations have emerged as a powerful tool for predicting O-glycoprotein structures, the high technical expertise and computational resources required limit the accessibility and throughput of these approaches. Recent efforts by the Fadda group have helped bridge these gaps through their development of GlycoShape, a publicly available modeling platform that enables rapid generation of glycoprotein models via a user-friendly web interface.^29^ Similarly, the Moremen group recently explored Alphafold 3.0 as a promising strategy for non-specialists to perform high-throughput structural prediction of glycoproteins and glycolipids.^30^ However, despite their efficiency and ease of implementation, these approaches do not readily capture measurable (glyco)protein features offered by traditional MD programs, such as conformational flexibility, hydrogen-bonding interactions, or solvent-accessible surface area.^31^ Notably, these properties account for protein behavior in a solvated, dynamic environment and therefore provide a more rigorous representation of protein structure than the static conformations generated by AlphaFold 3.0 and GlycoShape.

To address current challenges in studying lacritin glycosylation, we recently developed a glycoproteomics workflow to uncover the tear fluid glycoproteome, providing the first in-depth view of site-specific glycans which modify this protein.^16^ Here, we characterized highly diverse glycan structures, including the Tn antigen (GalNAcα1-Ser/Thr), sialylated core 1 (Galβ1-3GalNAcα1-Ser/Thr), type 3 H-antigen (Fuc α1-2-Galβ1-3GalNAcα1-Ser/Thr), and sialylated and fucosylated core 2 O-glycans (GlcNAcβ1-6(Galβ1-3)GalNAcα-Ser/Thr). In a follow-up study, we demonstrated that lacritin glycoforms harboring core 2 O-glycans could serve as a ligand for extracellular galectin-3 (Gal-3), a glycan-binding protein present in tear fluid.^32^ Interestingly, we also found that Gal-3 displayed higher affinity towards lacritin multimers compared to monomers, hinting at a potential difference in the glycosylation landscape between these two populations.

Herein, we hypothesized that multimeric and monomeric lacritin exhibit unique glycosylation profiles, and that these glycans could affect their structural conformation. More specifically, we asked whether glycans proximal to crosslinking residues (Lys101 and Lys104) could influence solvent accessibility at these sites, with potential implications for multimerization. To investigate this, we developed a sample preparation method which could separate lacritin multimers from monomers for downstream glycoproteomic characterization. From MS data, we quantified the most abundant glycoforms in each fraction and found differences in O-glycan heterogeneity at Ser86, Ser91, and Thr 95. Building on this, we leveraged publicly available software (CHARMM-GUI and GROMACS) to perform standard MD simulations on the most abundant glycoforms in our dataset. Here, computational analyses predicted that glycosylation could influence the flexibility of the two C-terminal alpha-helices through intra-glycan-protein interactions. Finally, we quantified the solvent-accessible surface area (SASA) of Lys101 and Lys104 and found that multimer-associated glycoforms exhibited significantly higher SASA than both monomeric and unmodified forms. Taken together, our study explores site-specific O-glycosylation as a key structural determinant of lacritin function, revealing a possible regulatory role for lacritin O-glycosylation. More broadly, the methods presented here are designed to be accessible to laboratories without specialized MD expertise and applicable to other O-glycosylated proteins.

## Results

### Development and validation of a method to characterize lacritin monomers and multimers

Given the distinct molecular weights of lacritin monomers (∼25 kDa) and multimers (≥ 50 kDa), we reasoned that GlycoFASP^33^ (filter-aided sample preparation) would provide an effective strategy for separating and characterizing these two populations. In brief, this workflow couples glycoprotease(s) with molecular-weight cutoff (MWCO) filters, which retain proteins above a designated MW while lower MW species are separated into the filtrate. We first collected tear fluid (6-8 μL collected by microcapillary) from three different healthy donors for immediate reduction and alkylation before loading onto a 30 kDa MWCO filter. After five rinses, monomers were recovered in the filtrate (<30 kDa) while multimers (>30 kDa) remained in the retentate (**Figure 1A, top**). Next, the filtrate was loaded onto a separate 10 kDa filter and both samples were processed identically using mucinase SmE to generate O-glycopeptides.^34,35^ Next, O-glycopeptides were collected through the filters and subjected to trypsin digestion and desalting before subsequent LC-MS/MS analysis (**Figure 1A, bottom**). Once MS raw files were generated, we manually validated MS spectra and performed label-free quantitation (LFQ) using area under the curve (AUC) intensities to quantify glycoforms found in each fraction (**Figure 1B, top**). This data was then used to inform downstream *in silico* analysis where AlphaFold 3.0 and CHARMM-GUI allowed us to build a glycosylated starting construct for MD simulations with GROMACS (**Figure 1B, bottom**). Finally, molecular structures and intra-glycan-protein interactions were visualized with Pymol.

**Figure 1.**
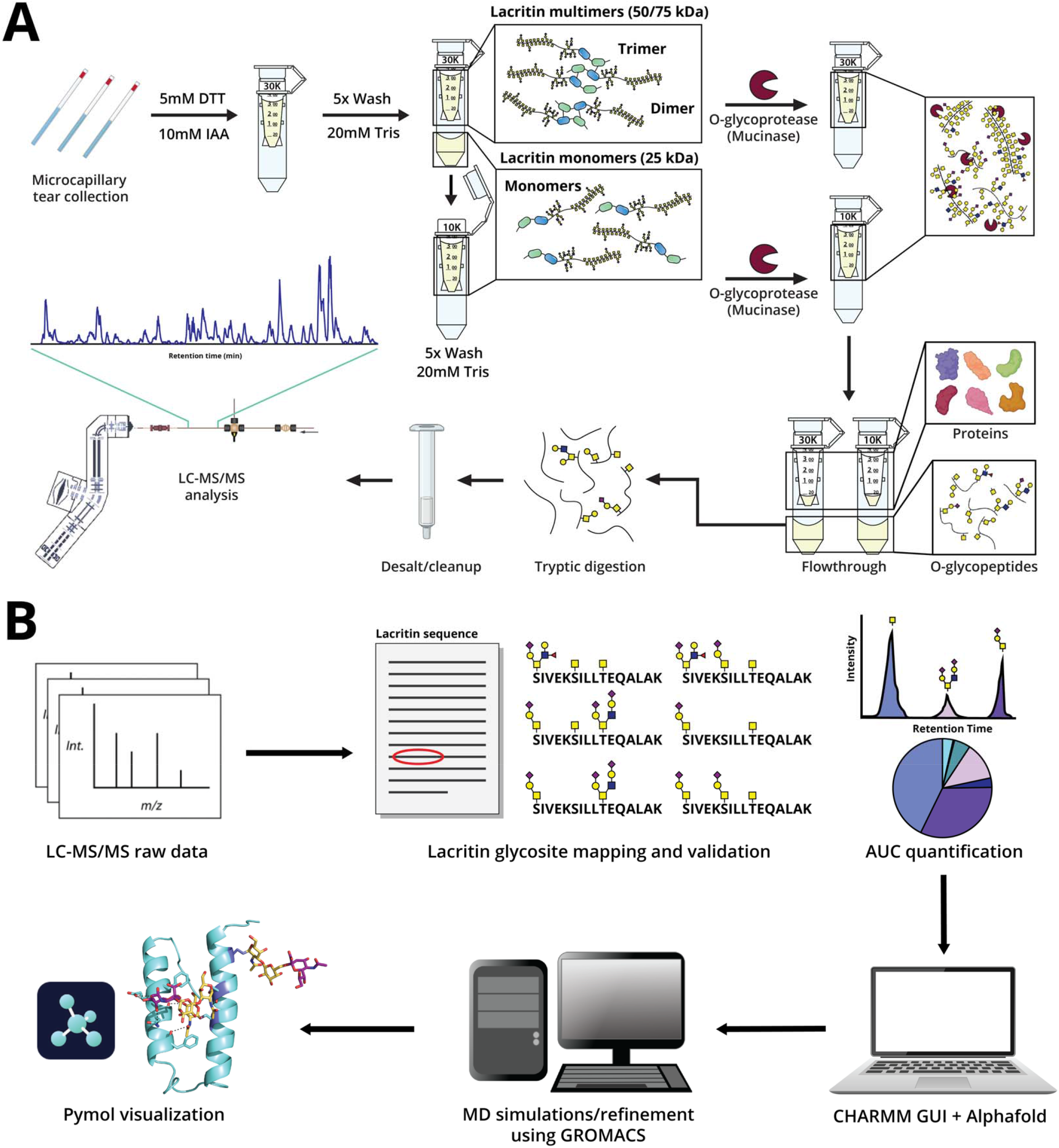
Glycoproteomic and molecular dynamics workflow to characterize lacritin. (**A**) Tear fluid (3-4 μL) was collected by microcapillary tubes and resuspended in 20mM Tris before being subjected to reduction and alkylation with dithiothreitol (DTT) and iodoacetamide (IAA). Next, the sample was washed 5 times on a 30 kDa MWCO filter (containing multimers) and the filtrate (containing monomers) was collected and loaded onto a separate 10 kDa MWCO filter. Next, O-glycoprotease (mucinase) SmE was added to the top of each filter to generate lacritin glycopeptides which are collected as the flowthrough. Finally, tryptic digestion was performed before desalting and downstream LC-MS/MS analysis. (**B**) MS raw files were searched with Byonic and lacritin glycopeptides were manually validated before quantitation with LFQ using AUC intensities. The abundances of validated glycopeptides were used to inform the starting glycoform construct which was built using CHARMM GUI and Alphafold 3.0. Lastly, molecular dynamics was executed with GROMACS and Pymol was used for molecular visualization.

Western blot analysis of control tear fluid (input) and the washed retentate across three biological replicates confirmed efficient separation of lacritin multimers from monomers, with the input containing both species and the retentate (after five washes) containing only multimeric lacritin (**Figure 2A**). While the filtrate was also collected for Western blot analysis, the concentration of lacritin collected after 5 sample rinses was too dilute to be detected by Western blot with our starting sample input. Nonetheless, we proceeded with downstream MS analysis and obtained near full sequence coverage of lacritin in both the 10 and 30 kDa preparations (**Figure 2B**). Further analysis of peptides generated from SmE and trypsin revealed that peptides containing Lys104 were only detectable in the monomer samples (highlighted pink in **Figure 2B**). Given that Lys104 is known to participate in TGM2-mediated crosslinking, tryptic cleavage at this site would likely be hindered or result in a crosslinked peptide fragment (as previously reported^15^) in the multimer fraction. As the crosslinked peptide was not detected by our methods, the lack of a tryptic peptide containing Lys104 in our multimer fraction likely suggested crosslinking and provided further confirmation that these two populations could be separated with our workflow.

**Figure 2.**
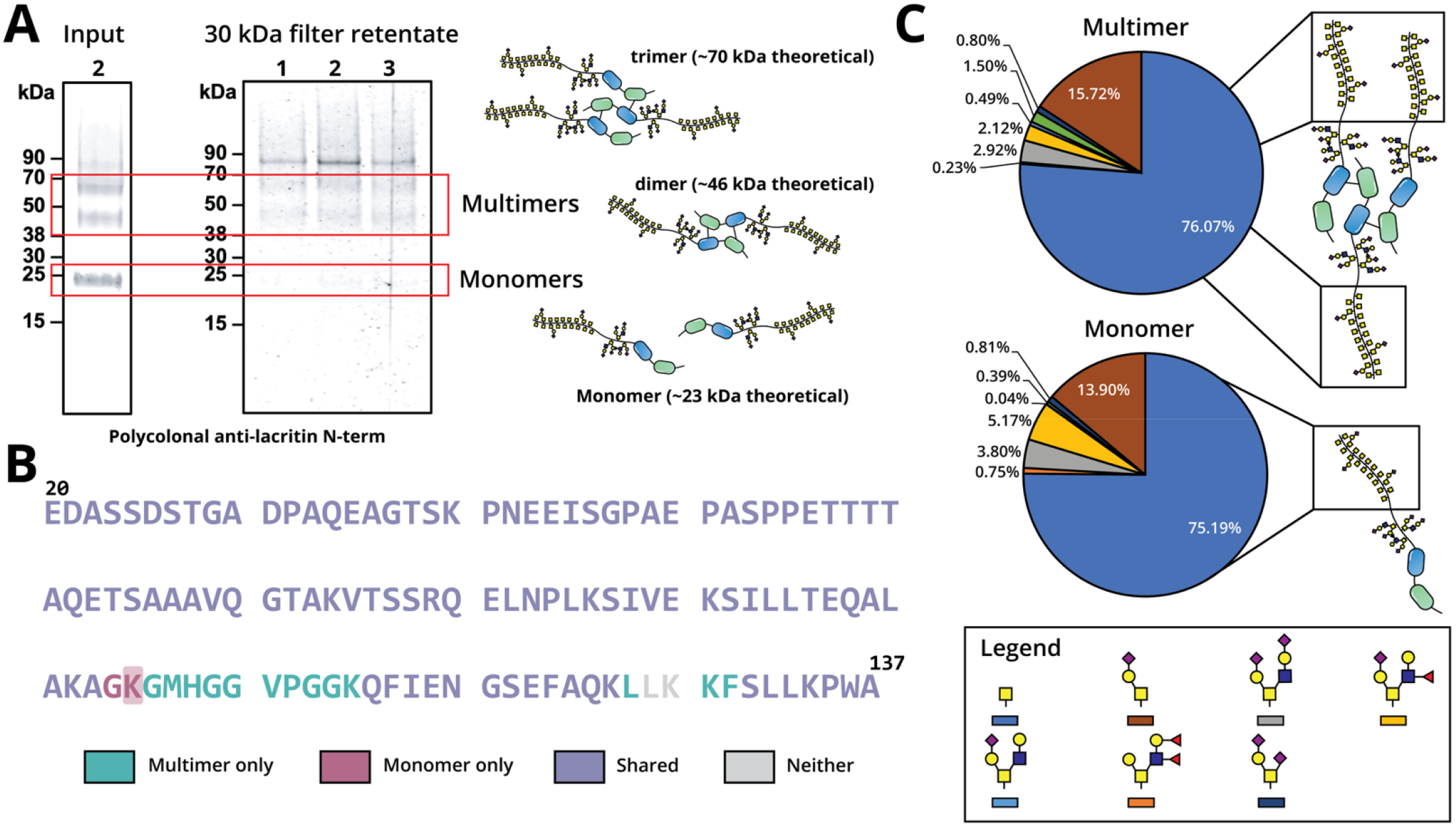
Validation of FASP workflow by Western blot and MS. (**A**) Western blot analysis of lacritin in tear fluid input (3-4 μL) and the 30kDa filter retentate after 5 washes using a polyclonal N-terminal anti-lacrtin antibody (‘anti-Pep Lac N-term’). (**B**) Lacritin sequence coverage map from downstream MS analysis of the monomer and multimer fraction after glycoFASP processing. Multimer sequence coverage is shown in teal, Monomer sequence coverage is shown in pink, and the shared sequence coverage is shown in purple. Coverage of one of the crosslinking lysines (Lys104) is highlighted in red and was identified only in the monomer fraction. (**C**) Quantification of N-terminal mucin domain glycosylation in both fractions. Glycan structures and their corresponding colors in the pie chart are denoted in the legend on the bottom.

Next, we characterized the glycosylation landscape of the N-terminal mucin domain (residues 23-77) in both fractions of a single patient. Interestingly, we observed highly similar glycan profiles with the Tn antigen as the most abundant structure detected (**Figure 2C, Table S1A**). Monosialylated core 1 and sialylated and fucosylated core 2 O-glycans were also detectable at lower relative abundances, consistent with our previous glycoproteomic analysis of lacritin.

### In-depth analysis of Ser86, Ser91, and Thr95 glycosylation

As previously noted, the C-terminal helices of lacritin participate in SDC1 binding, multimerization, and antimicrobial activity to modulate ocular homeostasis.^11,14,15^ Given the biochemical significance of this region and the difference in biological activity between monomers and multimers^15^, we hypothesized that site-specific O-glycans in the C-terminus (Ser86, Ser91, and Thr95) might differ between these two species. To investigate this, we first extracted ion chromatograms (XICs) for representative glycopeptides modified at these sites for both the monomer (**Figure S1A**) and multimer (**Figure S2A**) fractions across three different donors. To validate our replicates were comparable, we further examined glycopeptide retention times (**Figure S1B, Figure S2B**), coefficient of variation of the total lacritin intensity (**Figure S1C, Figure S2C**), and unique glycopeptide identifications (**Figure S2D, Table S1B**). Given that our samples were consistent throughout these analyses, we decided to focus on a single patient for in-depth quantitation using LFQ.

We assessed O-glycan heterogeneity at Ser86, Ser91, and Thr95 by compiling the AUC intensities of all glycopeptides spanning these glycosites (**Table S1C**). For Ser86, we observed that the monosialylated core 1 O-glycan was the most abundant for both populations, though multimers had a higher relative abundance of more diverse structures such as disialylated core 1 (10.88% vs. 3.60%) and core 2 O-glycans (19.62% vs. 3.17%) (**Figure 3A and B**). Interestingly, we observed that the multimer fraction displayed a higher relative abundance across all the core 2 structures found at this site. Most strikingly, we found that monomeric Ser91 was predominantly modified with a sialylated core 1 (83.95%), whereas in the multimer, Ser91 was largely unoccupied (42.77%) or decorated only by a single GalNAc (32.49%) (**Figure C and D**). Finally, Thr95 was primarily modified by the Tn antigen for both species (75.21% and 52.12%), though sialylated core 1 and core 2 O-glycans were also detectable (**Figure 3E and F**). These results highlight that the two lacritin populations exhibit distinct C-terminal glycosylation profiles, which could contribute to their differing biological roles. Although site-specific O-glycan differences have been reported for recombinant SARS-CoV-2 S monomer and dimer derived from insect cells, the extent to which O-glycan patterns coincide with multimeric state in native biological contexts remains underexplored. To the best of our knowledge, this study represents the first glycoproteomic observation of endogenous O-glycosylation variation between monomeric and multimeric species.

**Figure 3.**
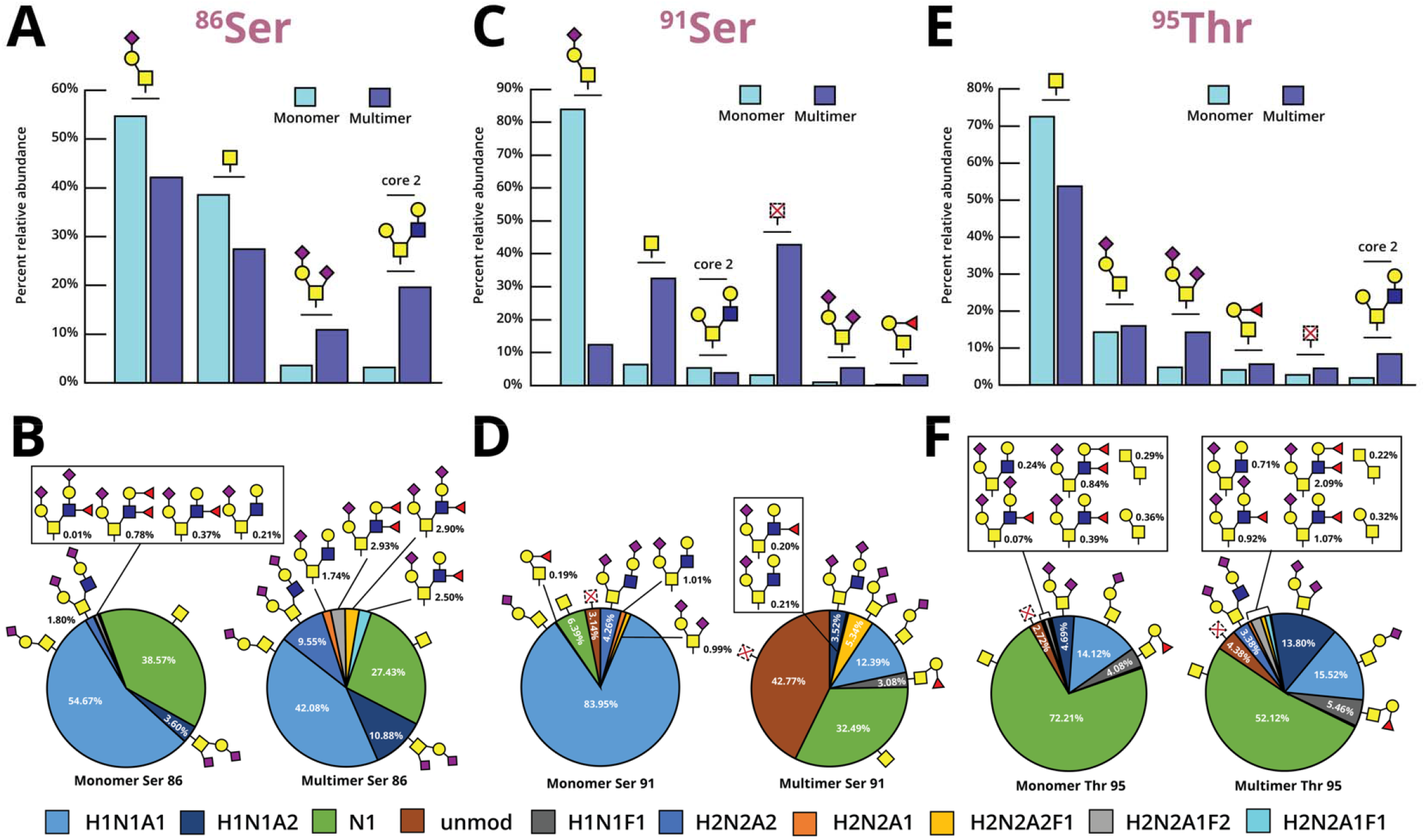
Site-specific O-glycoproteomic analysis and glycan quantitation at Ser86, Ser91, and Thr95. (**A**,**C**,**E**) All identified glycopeptides containing Ser86, Ser91, and Thr95 from a matched monomer and multimer fraction of a single patient was subjected to manual validation and LFQ using AUC intensities to generate corresponding bar graphs. Light blue bars correspond to the monomer and dark blue bars correspond to the multimer. (**B**,**D**,**F**) Pie chart representations of O-glycan heterogeneity at Ser86, Ser91, and Thrr95 in both fractions. In the colored legend below, H denotes Hexose, N denotes HexNAc, A denotes Neu5Ac, and F. denotes Fucose. Unmod describes no glycan occupancy at that site.

### Structural analysis of C-terminal lacritin with molecular dynamics

The presence of differential O-glycosylation within the ordered C-terminal domain prompted us to ask whether glycans might impart distinct conformational features. Notably, structural visualization of this region could provide insight into how O-glycans affect SDC1 binding and/or multimerization at the C-terminus of lacritin. To explore this, we modeled three C-terminal constructs (non-glycosylated, “monomer”, and “multimer”) spanning residues 79-137 of lacritin (**Figure 4A**). As described in **Figure 1B**, we used the most abundant glycans at Ser86, Ser91, and Thr95 (**Figure 3**) to represent the monomer and multimer glycoforms. Ser86 was modified with a monosialylated core 1 O-glycan and Thr95 was decorated with a single GalNAc for both species. At Ser91, a monosialylated core 1 O-glycan was used for the monomer while the multimer was left unmodified. To generate the initial structures for MD simulations, we leveraged AlphaFold 3.0 and CHARMM GUI which enabled us to build lacritin C-terminal constructs with defined glycosylation. Following this, we employed GROMACS for minimization, equilibration, and a 500 ns production step (see **Methods** for more detail).

**Figure 4.**
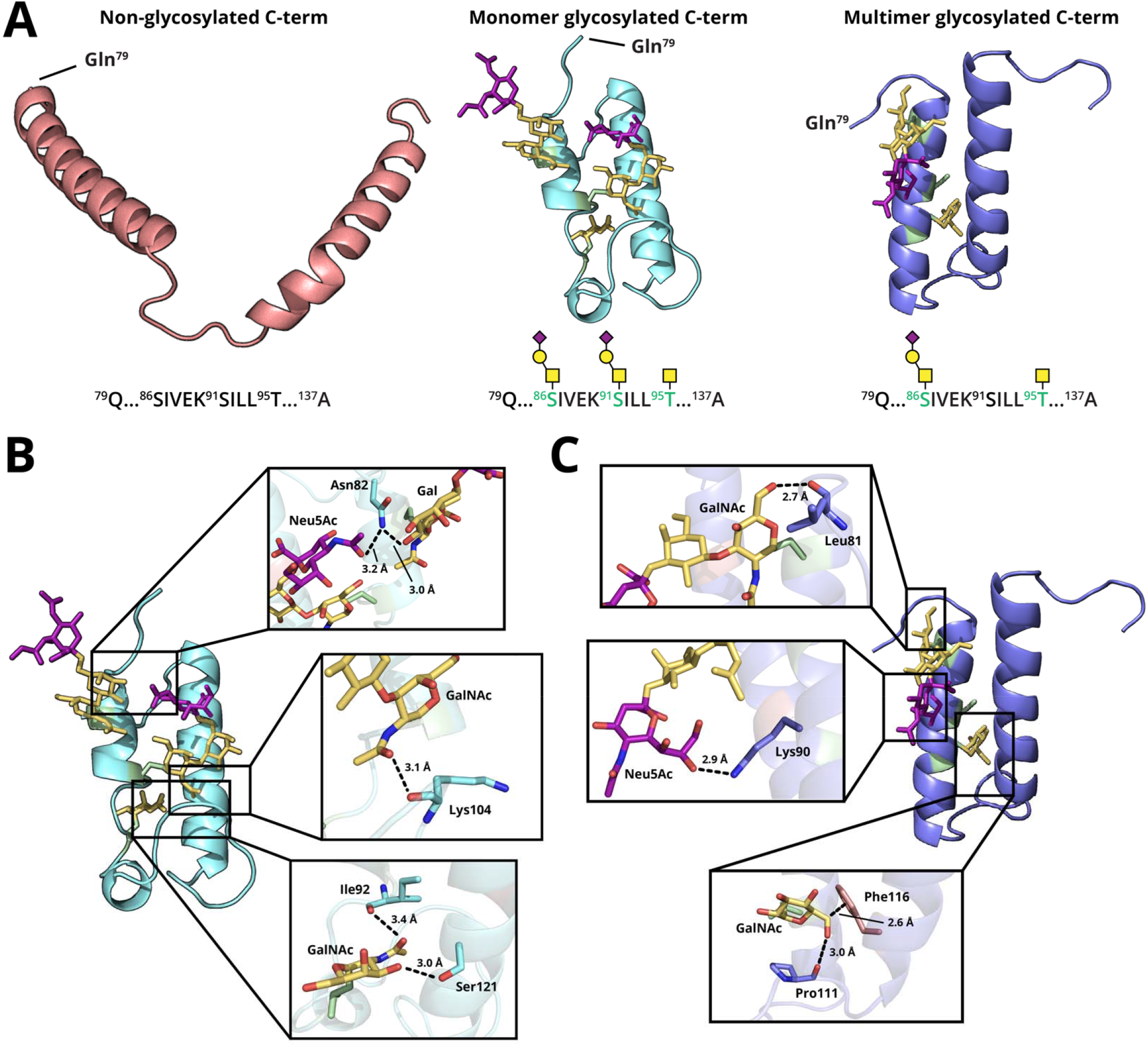
C-terminal lacritin structural analysis and glycan-protein interactions. (**A**) A 500 ns production step was executed on GROMACS for three C-terminal lacritin constructs (residues 79 to 137) and the 450 ns frame was extracted for molecular visualization. Below each structure is the sequence and corresponding glycan occupancy at Ser86, Ser91, and Thr95 as previously determined in **Figure 3** by taking the most abundant glycan structure at each site. The non-glycosylated structure, monomer-associated glycoform, and multimer-associated glycoform are colored in red, teal, and blue, respectively. (**B**) Intra-glycan-protein interactions (GPIs) in the monomer-associated glycoform are shown. Asn82 participates in polar contacts with Neu5Ac (3.2 Å) and Gal (3.0 Å) of Ser86 (top). The Lys104 backbone forms a polar contact with GalNAc (3.1 Å) of Ser91 (middle). Ser121 and Ile92 each make a polar contact with GalNAc (3.0 Å and 3.4 Å) of Thr95 (bottom). (**C**) Intra-GPIs in the multimer-associated glycoform are depicted where the Leu81 backbone has a polar contact with GalNAc (2.7 Å) of Ser86 (top). Lys90 forms a polar contact with Neu5Ac (2.9 Å) of Ser86 (middle). Pro111 makes a polar contact with GalNAc (3.0 Å) of Thr95 while Phe116 participates in pi-stacking (2.6 Å) with the same GalNAc (bottom). PDB files of the extracted frame were visualized on Pymol and distances of polar contacts were traced on Pymol. The final structure was rendered and exported onto adobe illustrator for residue and glycan labeling.

The resultant structures shown in **Figure 4A** represent frames extracted at the 450 ns time point, which was used for subsequent structural analysis. Based on these structures, we hypothesized that the proximity and positioning of glycans relative to nearby amino acids could reveal intra-glycan-protein interactions (GPIs) across the two alpha helices. These polar contacts could influence the local conformation and positioning of the two helices, which may have implications for downstream binding events. Using the predominant glycoform of the monomer, we showed that Asn82 can participate in hydrogen-bond interactions with the C2 hydroxyl on Gal of Ser86 (3.0 Å) and the N-acetyl on Neu5Ac of Ser91 (3.2 Å). Additionally, the GalNAc N-acetyl on Ser91 can coordinate with the Lys104 peptide backbone (3.1 Å) while Ile92 and Ser121 participate in polar interactions with the N-acetyl and C3 hydroxyl of GalNAc on Thr95 (**Figure 4B**). Interestingly, these specific contacts were not present in the frame extracted for the multimer glycoform. Instead, we observed an altered positioning of the Ser86 glycan driven by hydrogen-bond interactions between the Leu81 peptide backbone and the GalNAc C6 hydroxyl (2.7 Å) as well as between Lys90 and the Neu5Ac C8 hydroxyl (2.9 Å). In the flexible loop connecting the two helices, Pro111 formed dipole interactions with the Thr95 GalNAc C6 hydroxyl (3.0 Å), which is partially held in place by a potential pi-stacking interaction with Phe116 (∼2.6 Å) (**Figure 4C**). Lastly, we tracked the frequency of intra-glycan-protein hydrogen-bond interactions for both constructs and found a similar number between replicates (**Figure S3, Table S1D**). Overall, these results highlight a possible role for site-specific O-glycans in affecting the structural topology of C-terminal lacritin through intra-protein polar contacts. Specifically, differences in glycan heterogeneity are predicted to result in distinct residue-glycan interactions, potentially affecting the spatial arrangement of the protein backbone. As such, we envision that these models may inform hypotheses surrounding lacritin C-terminal binding interactions or protease processing of this domain.

### O-glycans affect C-terminal flexibility and SASA of crosslinking residues

Given the prevalence of intra-GPIs in our glycosylated constructs, we hypothesized that the overall flexibility of the protein backbone could be influenced by these interactions. In particular, intra-GPIs may help to localize the two helices closer together and promote a more “rigid” structure. In contrast, removal of glycans is predicted to result in substantially decreased electrostatic interactions that stabilize the two helices, leading to increased conformational flexibility (**Figure 5A**). To support this reasoning, we quantified the flexibility of the protein backbone (as measured by root mean square deviation-RMSD) for all three constructs over the 500 ns production step. Here, we showed that the unmodified construct was significantly more flexible over the entire duration of the simulation, with an average RMSD of ∼1.4 nm compared to ∼0.6 nm when glycosylated (**Figure 5B, Table S1E**). This measurement was validated across three replica simulations with randomized initial velocities, where the average RMSD and standard error is plotted for each structure (**Figure 5C, Table S1E**). Overall, this data indicated that O-glycans are predicted to alter the conformational flexibility of C-terminal lacritin when compared to the unmodified structure. More broadly, our observation that O-glycans have projected effects on the rigidity of the protein backbone has been supported in previous MD studies^36,37^ and by atomic force microscopy.^28,38^

**Figure 5.**
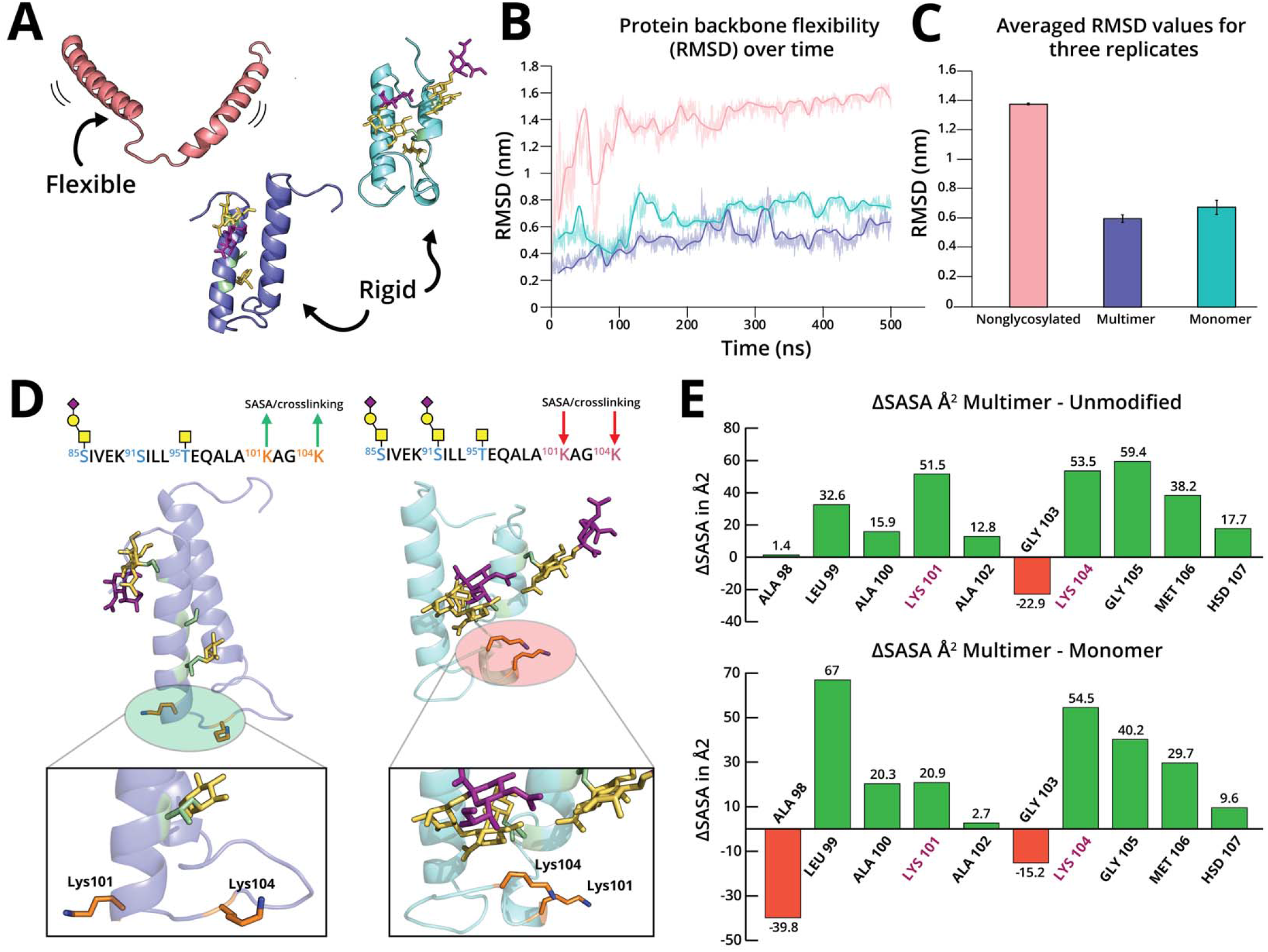
C-terminal O-glycans affect protein flexibility and the SASA of crosslinking residues. (**A**) Protein structures after MD simulations were labeled with a qualitative description of protein flexibility. (**B**) Protein backbone flexibility (RMSD in nm) was measured for a single replica over the 500ns production step for the nonglycosylated (pink), “monomer” (teal), and “multimer” (dark blue) glycoforms. (**C**) Three independent replica simulations for each construct was executed, and the average RMSD over the 500ns production step was taken for each for a total of 9 data points. The bar graph shows the average RMSD value for the three structures, where each bar represents three replicate simulations. (**D**) Molecular visualization of multimer and monomer-associated glycoforms and the impact of glycosylation on Lys101 and Lys104 orientation (using the 450ns frame). In the monomer structure, glycans point in the same direction as Lys 101 and 104 and are closer in proximity to the lysines compared to the multimer glycoform. (**E**) The average solvent accessible surface area (SASA) in Å^2^ was taken across a window spanning three residues upstream of Lys101 to three residues downstream of Lys104 for each construct. Next, the difference in SASA (ΔSASA) was taken between the Multimer and the unmodified structure (top) and between the Multimer and Monomer (bottom). Green bars represent a predicted increase in SASA while red bars represent a predicted decrease. See the **Methods** section and **Table S1E-G** for more details.

Building on these findings, we asked whether specific glycoforms could provide structural insights into propensity for multimerization, where residues that participate in crosslinking might be affected by proximal glycans. As shown in **Figure 5D (left structure)**, Lys101 and Lys104 of the multimer-associated glycoform are anticipated to position in solvent exposed areas, which may promote access to TGM2 for multimerization. Alternatively, monosialylated core 1 O-glycans on the monomer glycoform (**Figure 5D, right structure**) are positioned in closer proximity to Lys101 and Lys104 through hydrogen-bonding interactions with the Lys104 backbone and orient in the same direction as these residues. In this configuration, Lys101 and Lys104 appear less solvent accessible, with additional steric effects potentially imposed by the sialylated core 1 O-glycans due to their spatial orientation. To justify these qualitative observations, we extracted the average SASA (per residue) and quantified their difference (ΔSASA) between our modeled constructs. By calculating the ΔSASA across a window spanning three residues upstream of Lys101 to three residues downstream of Lys104, we observed a general increase in solvent accessibility for the multimer-associated glycoform relative to the monomeric and non-glycosylated structures (**Figure 5E, Table S1F**). Most notably, the SASA of Lys101 and 104 were predicted to increase by 51.5 Å^2^ and 53.5 Å^2^, respectively, when comparing the multimer to the unmodified C-terminus. Similarly, the SASA of Lys101 and 104 were predicted to increase by 20.9 Å^2^ and 54.5 Å^2^, respectively, when comparing multimer and monomer glycoforms. These trends were recapitulated across three replica simulations, where the averaged ΔSASA at Lys101 and 104 were +14.97 Å^2^ and +40.83 Å^2^, respectively, when comparing the two glycoforms (**Table S1G**). Collectively, these analyses demonstrate that there are predicted structural differences imparted by site-specific O-glycans, with implications for affecting lacritin multimerization.

## Discussion

To date, the functional role of lacritin O-glycosylation has remained a critical blind spot in our understanding of its biology. To be sure, the chemical complexity of O-glycan heterogeneity has largely impaired biochemical, bioanalytical, and structural analyses of lacritin glycoforms over the past three decades. In our prior study^16^, we leveraged recent advances in MS-based glycoproteomics^35,37,39,40^ to characterize the tear fluid glycoproteome, establishing the first in-depth view of site-specific O-glycans which decorate lacritin. Subsequent analysis with AlphaFold 3.0 predicted that O-glycosylation imparts structural rigidity and a “bottlebrush-like” secondary structure, in line with observations for other mucin-domain glycoproteins.^16,36^ From additional follow-up studies^18,32^, we demonstrated that the densely O-glycosylated region of lacritin can confer proteolytic protection or serve as a ligand for Galectin-3 (Gal-3) in tear fluid. In this context, the mucin domain may modulate the extracellular half-life of lacritin by sterically blocking protease access through its O-glycans. Alternatively, engagement of Gal-3 as an extracellular binding partner may likewise influence circulating half-life, consistent with observations reported for other extracellular glycoproteins.^41,42^

Building on these insights, we hypothesized in the present study that O-glycans affect the C-terminal domain of lacritin, which has implications for its downstream biological activity and ocular homeostasis. Notably, multimerization stands out as a key regulatory mechanism which mediates lacritin effector function, since multimeric lacritin is incapable of binding SDC-1 to trigger GPCR signaling. This led us to ask whether multimers and monomers differed in their O-glycosylation patterns, particularly at glycosites proximal to crosslinking residues (Lys101 and Lys104). To investigate this, we modified our previously reported GlycoFASP method to isolate lacritin multimers and monomers for separate downstream MS analyses. Here, we discovered distinct glycoform distributions in each fraction and subsequently quantified site-specific glycan abundances to inform the design of C-terminal lacritin constructs for MD simulations.

From *in silico* analyses, O-glycans were predicted to participate in intra-GPIs between the two alpha helices, positioning them closer to each other via electrostatic interactions. We expanded on these observations by measuring protein flexibility and revealed that the protein backbone was projected to be more flexible without glycosylation, which has also been shown for other glycosylated proteins.^36,37,43^ Finally, we used basic MD analysis tools to evaluate the SASA of Lys101 and 104, demonstrating that these residues were highly solvent exposed in the multimer-associated glycoform, whereas non-glycosylated and monomeric glycoforms displayed lower SASA values at these sites. Future comparison of the *in vitro* crosslinking efficiencies of different lacritin constructs with defined glycosylation patterns will be informative for validating the structural observations predicted in our analyses, although such experiments are not yet feasible with current strategies for recombinant lacritin expression. Nonetheless, these results point to a structural role for O-glycosylation in influencing multimerization, where intra-GPIs may play a key function. Consistent with this, a study by Wu and Robinson showed that trimeric tumor necrosis factor-⍰ (TNF-⍰) is stabilized by an O-linked glycan at Ser80, while Karampini et al. demonstrated that von Willibrand factor multimerization is altered upon treatment of endothelial cells with an O-glycosylation inhibitor (GalNAc-O-benzyl).^44,45^ Although these studies highlight an emerging role for O-glycan-driven effects on multimerization, this phenomenon remains largely underexplored, with only these few examples reported to date. Our findings expand on the growing body of literature on O-glycan-mediated multimerization and underscore the importance of discovering these interactions in other O-glycoproteins.

Taken together, this study lays the critical groundwork for future investigation of lacritin biology mediated by its glycans. It currently remains unknown whether distinct lacritin glycoforms might exhibit varying affinities towards SDC1 or engage different protein ligands entirely. Similarly, the extent to which glycans might affect the antimicrobial activity of C-terminal lacritin peptides will require future studies to elucidate. Finally, spliceoform-specific multimerization has also been reported^46^, where isoform C and (potentially) isoform D can also multimerize. Whether the canonical splice variant forms heterodimers with isoform C or D remains unclear, though it is likely that O-glycans would play a role in this process as well. As glycosylation vastly expands the chemical space of lacritin, it will be necessary to account for this additional layer of complexity when defining new roles for lacritin at the ocular surface. From a diagnostic perspective, it is possible that lacritin glycosylation could be concomitant with ocular pathologies such as DED. Though it has not been established whether lacritin monomers or multimers offer more diagnostic potential based on their disease-associated glycan changes, the methods described in this study now enable these two populations to be separated for independent glycoproteomic characterization. Further efforts to profile their glycosylation changes across ocular diseases will thus be crucial to inform glycan-centric diagnostic strategies.

More broadly, this study highlights MS-based O-glycoproteomics and basic MD simulations as complementary analytical tools to explore fundamental biochemistry mediated by glycans. Until recently, the ability to identify and quantify O-glycans in a site-specific manner with high sensitivity has presented an enormous analytical challenge. This is evidenced by recent studies where MD modeling of O-glycoproteins rely on released O-glycan analysis from large quantities of a purified glycoprotein to obtain the glycan data needed to inform simulations.^47,48^ Though glycoproteomics experiments can be performed as an alternative to obtain site-specific O-glycan data, accurate and sensitive characterization of low abundance glycoproteins from complex samples remains a critical bottleneck in the glycoproteomics workflow.^49–51^ To address these limitations, recent advances in MS-based glycoproteomics such as improved enrichment strategies^52^, optimized gas-phase fragmentation methods^53^, and the use of O-glycoproteases and mucinases have enhanced our ability to map O-glycosylated proteins.^40,54^ This is highlighted in a recent collaboration between our laboratory and the Amaro group^37^, where we demonstrated that site-specific glycoproteomics data can be coupled with MD simulations to achieve detailed models of densely O-glycosylated mucins at the cell surface. However, the technical expertise required, reliance on specialized in-house MD software, and substantial computational demands (several months on a supercomputer) decrease the accessibility of this approach to non-specialists and laboratories without MD capabilities. As such, looking forward, we envision that the methods described in this study will provide an accessible framework for integrating high quality O-glycoproteomics data with publicly available MD software to investigate structural effects driven by site-specific O-glycans.

## Materials and Methods

### Tear fluid Sample collection

Tear fluid was purchased from Innovative Research (obtained via microcapillary collection) and stored at −20°C until further processing. 3 individual donors (56669-ND0557-CF35, 56668-ND0350-CF42, and 56667-ND0143-CF43) were used for western blot and downstream MS analysis. According to the vendor’s collection documentation, all tears were collected under basal (non-stimulated) conditions, and no products or procedures were used to stimulate tear production (no local anesthesia was used). All donors were classified by the vendor as “normal donors” after passing an ocular history and overall health risk-factor screening questionnaire. Patients also provided signed clearance for collection. The ocular screening questionnaire evaluates any current or recent history of dry eye disease, eye irritation, ocular surgery, current redness or infection or inflammation, contact lens use and recency of wear, use of ocular medications/eye drops, allergy history and recent active allergic symptoms. Screening of general health status evaluated recent infection, chronic medical conditions, pregnancy/lactation status, prescription medication use (including immunosuppressants/steroids/ biologics), recent antibiotic/antiviral use, infectious disease history and exposure risk (including HIV, HBV, HCV), recent transfusion, recent tattoos/piercings, recent active allergic symptoms, smoking/vaping status, recent alcohol intake, recent exposure to air pollution/chemicals/smoke, recent illness, recent vaccination, and recent COVID-19 infection. See **Table S1H** for more details.

### Mass spectrometry sample preparation with GlycoFASP

Freshly thawed tear fluid proteins from each patient were diluted to a final concentration of 0.2 mg/mL in 100 µL of 20 mM Tris. DTT was then added to a concentration of 2 mM and reacted at 65 °C for 1 hour followed by alkylation in 5 mM IAA for 15 min in the dark at RT. Subsequently, samples were loaded onto a 30 kDa filter and the filtrate was collected over five washes (400 µL of 20mM Tris for each elution). The filtrate was then loaded onto a 10 kDa filter and denoted as the “monomer fraction” whereas the retentate of the 30 kDa filter is the “multimer fraction”. For both fractions, mucinase SmE was added at a 1:3 (E:S) ratio and allowed to react for 12 hours at 37 °C. After glycoprotease digestion, O-glycopeptides were filtered into a separate tube and subjected to an additional trypsin digestion for 3 hours at 37 °C. Finally, peptides were acidified by adding 4 µL of formic acid and desalted before injection into the mass spectrometer. Desalting was performed using 10 mg Strata-X 33 µm polymeric reversed phase SPE columns (Phenomenex). Each column was activated using 500 µL of acetonitrile (ACN) (Honeywell) followed by of 500 µL of 0.1% formic acid, 500 µL of 0.1% formic acid in 40% ACN, and equilibration with two additions of 500 µL of 0.1% formic acid. After equilibration, the samples were added to the column and rinsed twice with 200 µL of 0.1% formic acid. The columns were transferred to a 1.5 mL tube for elution by two additions of 150 µL of 0.1% formic acid in 40% ACN. The eluent was then dried using a vacuum concentrator (LabConco) prior to reconstitution in 10 µL of 0.1% formic acid. The resultant peptides were then injected onto a Dionex Ultimate3000 coupled to a Thermo Orbitrap Eclipse Tribrid mass spectrometer. We employed a higher-energy collision dissociation product-dependent electron transfer dissociation (HCD-pd-ETD) method; in some cases, we used supplemental activation in ETD (EThcD). The files were searched using Byonic, followed by manual data curation.

### Mass spectrometry data acquisition

Samples were analyzed by online nanoflow liquid chromatography-tandem mass spectrometry using an Orbitrap Eclipse Tribrid mass spectrometer (Thermo Fisher Scientific) coupled to a Dionex UltiMate 3000 HPLC (Thermo Fisher Scientific). For each analysis, 4 µL was injected onto an Acclaim PepMap 100 column packed with 2 cm of 5 µm C18 material (Thermo Fisher, 164564) using 0.1% formic acid in water (solvent A). Peptides were then separated on a 15 cm PepMap RSLC EASY-Spray C18 column packed with 2 µm C18 material (Thermo Fisher, ES904) using a gradient from 0-35% solvent B (0.1% formic acid with 80% acetonitrile) in 60 min. Full scan MS1 spectra were collected at a resolution of 60,000, an automatic gain control target of 3e5, and a mass range from *m/z* 300 to 1500. Dynamic exclusion was enabled with a repeat count of 2, repeat duration of 7 s, and exclusion duration of 7 s. Only charge states 2 to 6 were selected for fragmentation. MS2s were generated at top speed for 3 seconds. Higher-energy collisional dissociation (HCD) was performed on all selected precursor masses with the following parameters: isolation window of 2 m/z, 29% normalized collision energy, orbitrap detection (resolution of 7,500), maximum inject time of 50 ms, and a standard automatic gain control target. An additional electron transfer dissociation (ETD) fragmentation of the same precursor was triggered if 1) the precursor mass was between m*/z* 300 to 1500 and 2) 3 of 8 HexNAc or NeuAc fingerprint ions (126.055, 138.055, 144.07, 168.065, 186.076, 204.086, 274.092, and 292.103) were present at *m/z* ± 0.1 and greater than 5% relative intensity. Two files were collected for each sample: the first collected an ETD scan with supplemental energy (EThcD) while the second method collected a scan without supplemental energy. Both used charge-calibrated ETD reaction times, 100 ms maximum injection time, and standard injection targets. EThcD parameters were as follows: Orbitrap detection (resolution 7,500), calibrated charge-dependent ETD times, 15% nCE for HCD, maximum inject time of 150 ms, and a standard precursor injection target. For the second file, dependent scans were only triggered for precursors below *m/z* 1000, and data were collected in the ion trap using a normal scan rate.

### Mass spectrometry data analysis

Raw files were searched using Byonic (version 4.5.2, Protein Metrics, Inc.) against a FASTA file containing the canonical lacritin sequence (acession number: Q9GZZ8) and its spliceoforms (acession numbers: H0Y100 and F8W0V3). For all samples, we used the default O-glycan database containing 9 common structures. Files were searched with six missed cleavages and N-terminal to Ser/Thr and C-terminal to Arg/Lys. Mass tolerance was set to 10 ppm for MS1’s and 20 ppm for MS2’s. Pyroglutamylation and deamidation was set as a variable modification and carbamidomethyl Cys was set as a fixed modification. From the Byonic search results, glycopeptides were filtered to a score of >200 and a logprob of >2. From the remaining list of glycopeptides, the extracted ion chromatograms, full mass spectra (MS1s), and fragmentation spectra (MS2s) were investigated in XCalibur QualBrowser (Thermo) to generate a list of true-positive glycopeptides, as reported in **Table 2**. Each reported glycopeptide listed in **Table 2** was manually validated from the filtered list of Byonic’s reported peptides (score>200 and logprob >2) according to the following steps: The MS1 was first used to confirm the precursor mass and chosen isotope was correct. This also allowed us to identify any co-isolated species that could interfere with the MS2s and/or explain unassigned peaks. The HCD and EThcD fragmentation spectra were then investigated to identify sufficient coverage to make a sequence assignment. When possible, multiple MS2 scans were averaged to obtain a stronger spectrum. For HCD, an initial glycopeptide identification was confirmed if the presence of the precursor mass without a glycan present (i.e., Y0), along with coverage of b and y ions without glycosylation. For longer peptides, we required the presence of Y0 and fragments that were expected to be abundant (e.g., N-terminally to Pro, C-terminally to Asp). When the peptide contained a Pro at the C-terminus, the b_n-1_ was considered sufficient. Further, when the sequence contained oxidized Met, the Met loss from the bare mass was considered as representative of the naked peptide mass. We then used electron-based fragmentation MS2 spectra for localization. Here, all plausible localizations were considered, regardless of search result output. We confirmed the presence of fragment ions in ETD or EThcD that were between potential glycosylation sites, if sufficient c/z ions were present then a glycan mass was considered localized. For glycopeptide manual validation, extracted ion chromatograms are evaluated at the MS1 level to determine the charge and m/z of the highest abundance precursor species. Mass spectrometry data files and raw search output can be found on PRIDE with identifier PXD076054.

### MD analysis of lacritin C-terminal constructs

The most abundant O-glycan structures of lacritin were first calculated at each O-glycosite using LFQ of AUC intensities of XICs of all lacritin glycopeptides identified. Once glycan structures were determined, the sequence of C-terminal lacritin (Uniprot ID:Q9GZZ8, residues 77-137) was used to generate a pdb file on AlphaFold 3.0 webserver. The “nonglycosylated construct” consisted of the base C-terminal sequence without glycans. The “glycosylated construct” was generated by using 3-letter Chemical Component Dictionary (CCD) codes for glycans which were manually input onto Ser/Thr. Here, NAG was used for a single HexNAc residue. The resultant PDB file with the predicted fold was input into CHARMM-GUI for further glycan editing and to obtain a solvated and electrically neutral system in a rectangular periodic boundary water box. KCl ions were added via the Monte Carlo method at a concentration of 150 mM. CHARMM-GUI output files (.pdb solvated structure files, carbohydrate_restraint.str files, equilibration.inp and production.inp files) were then generated for downstream GROMACS analysis. Following system construction, initial geometry refinement was carried out in GROMACS using the steepest-descent algorithm for 5,000 steps, employing a maximum force tolerance of 1,000 kJ/mol·nm^−1^. The minimized structures were then equilibrated using the CHARMM-GUI multistage protocol, consisting of NVT heating and NPT equilibration at 303.15 °K and 1 atm. All equilibration steps were run with a 1 fs integration time step. The final equilibrated configuration (step4.1_equilibration.gro and step4.1_equilibration.cpt) served as the starting point for the production simulations. 500ns production simulations were performed in the NPT ensemble at 303.15 °K and 1 atm on an NVIDIA A5000 GPU node of the Yale Grace high-performance computing cluster. Following production, solvent-accessible surface area (SASA) and RMSD calculations were performed using GROMACS analysis tools (gmx rmsf, gmx sasa). To ensure adequate sampling of conformational space, two additional production replicas were performed for each construct. Replicas were initiated from the same equilibrated structure but with independently randomized initial velocities generated immediately after the equilibration stage. All replicas were run under identical thermostat, barostat, and integration settings. In total, three 500-ns production trajectories were generated per construct, yielding 1.5 μs of aggregate sampling. Replica consistency was assessed by comparing RMSD traces, SASA distributions, and qualitative inspection of alignment overlays across independent trajectories.

## Supporting information

This article contains supporting information.

## Data availability statement

All mass spectrometry data and search results acquired for this manuscript have been deposited on the PRIDE repository. Reviewers can access raw data with the following login information: Project accession: PXD076054 Token: qrOsmudfTPhF Alternatively, reviewer can access the dataset by logging in to the PRIDE website using the following account details:

Username: Password: 7mhpq1TvRHbN

## Author contributions

V.C. and S.A.M. conceptualization; V.C., R.J.C., and I.L. data curation; V.C., R.J.C., and I.L formal analysis; V.C. and R.J.C. investigation; V.C., R.J.C, and K.E.M. methodology; V.C. validation; V.C., R.J.C., and I.L. visualization; V.C. and S.A.M. writing– original draft; V.C., R.J.C, and S.A.M. writing–review & editing; S.A.M. and K.E.M. supervision; S.A.M. funding Acquisition; V.C. project administration; S.A.M. resources.

## Funding and additional information

V.C. is supported by an NSF GRFP (DGE-2139841).S.A.M. is supported by CRI Lloyd J Old STAR Award and a NIGMS R35-GM147039. I.L. is supported by The Yale College First-Year Summer Research Fellowship in the Sciences and Engineering.

## Abbreviations

The abbreviations used are:

ACN: Acetonitrile
ETD: Electron transfer dissociation
EThcD: Electron transfer/higher-energy collision dissociation
HCD: Higher-energy collisional dissociation
DED: Dry eye disease
TGM2: Transglutaminase 2
Tn antigen: GalNAcα1-Ser/Thr
sialylated core 1: Galβ1-3GalNAcα1-Ser/Thr
type 3 H-antigen: Fuc α1-2-Galβ1-3GalNAcα1-Ser/Thr
core 2: (GlcNAcβ1-6(Galβ1-3)GalNAcα-Ser/Thr;

