## Supplemental Information for "Site-specific O-glycans influence lacritin structure and multimerization in tears"

**Supplementary Table**

Supplementary Table S1A: Identified Lacritin mucin domain glycopeptides
Supplementary Table S1B: CV, AUC, RT, UGP analysis
Supplementary Table S1C: Ser86,Ser91, and Thr95 glycosylation
Supplementary Table S1D: Intra-glycan-protein H-bonds over time
Supplementary Table S1E: RMSD calculations
Supplementary Table S1F: SASA values
Supplementary Table S1G: Lys101 and Lys104 SASA replicate analysis

**Supplementary Figures**

Supplementary Figure 1. Lacritin monomer replicate analysis
Supplementary Figure 2. Lacritin multimer replicate analysis
Supplementary Figure 3. Intra-glycan-protein H-bonds over time

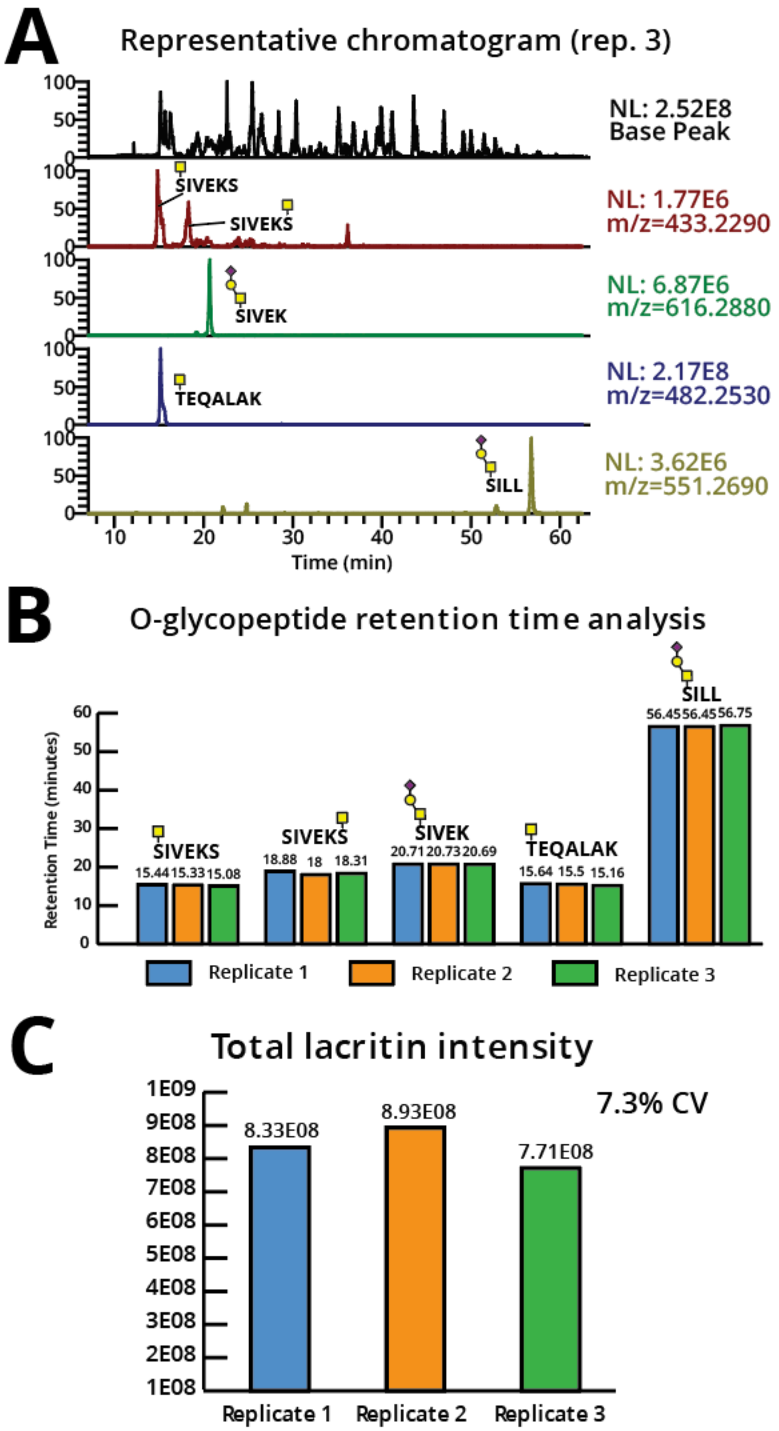


**Supplementary Figure 1. Lacritin monomer replicate analysis.** Tears from three healthy patients were collected by microcapillary tubes and processed separately from each other using the previously described glycoFASP method. (**A**) A representative extracted ion chromatogram of glycopeptides found in the monomeric lacritin fraction of replicate 3. (**B**) Retention time analysis of the glycopeptides extracted in (**A**) for each biological replicate. (**C**) Total intensity of lacritin monomer for each replicate, where %CV is caclulated using the formula: (standard deviation of intensity / mean intensity ) x 100%.

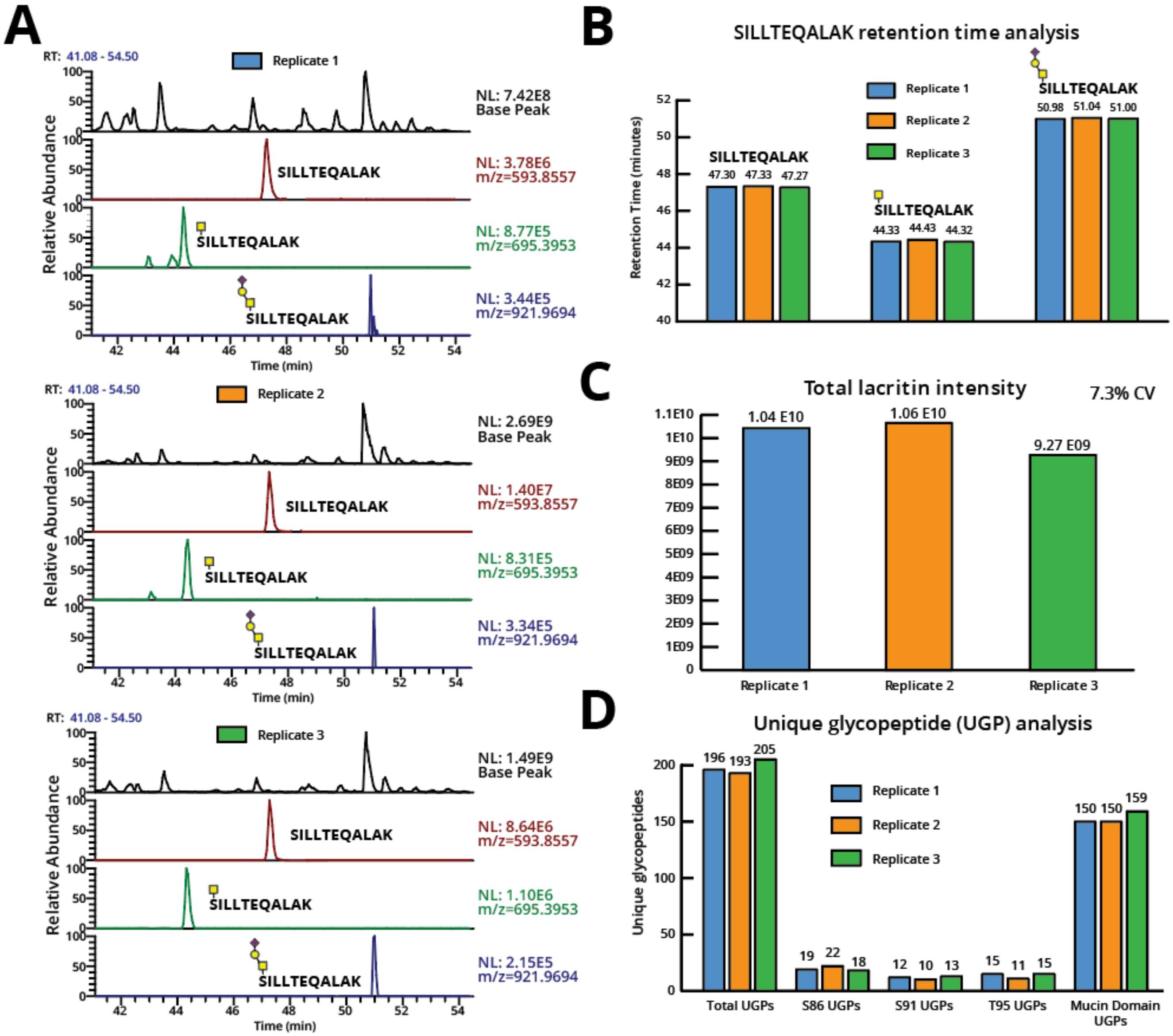


**Supplementary Figure 2. Lacritin multimer replicate analysis.** Tears from three healthy patients were collected by microcapillary tubes and processed separately from each other using the previously described glycoFASP method. (**A**) Extracted ion chromatograms for respresentative (glyco)peptides found in the multimeric lacrtin fraction across three biological replicates. (**B**) Retention time analysis of the glycopeptides extracted in (**A**) for each biological replicate. (**C**) Total intensity of lacritin multimer for each replicate, where %CV is caclulated using the formula: (standard deviation of intensity / mean intensity ) x 100%. (**D**) Unique glycopeptides (UGPs) identified for Ser 86, Ser 91, Thr 95, and the mucin domain for each biological replicate.

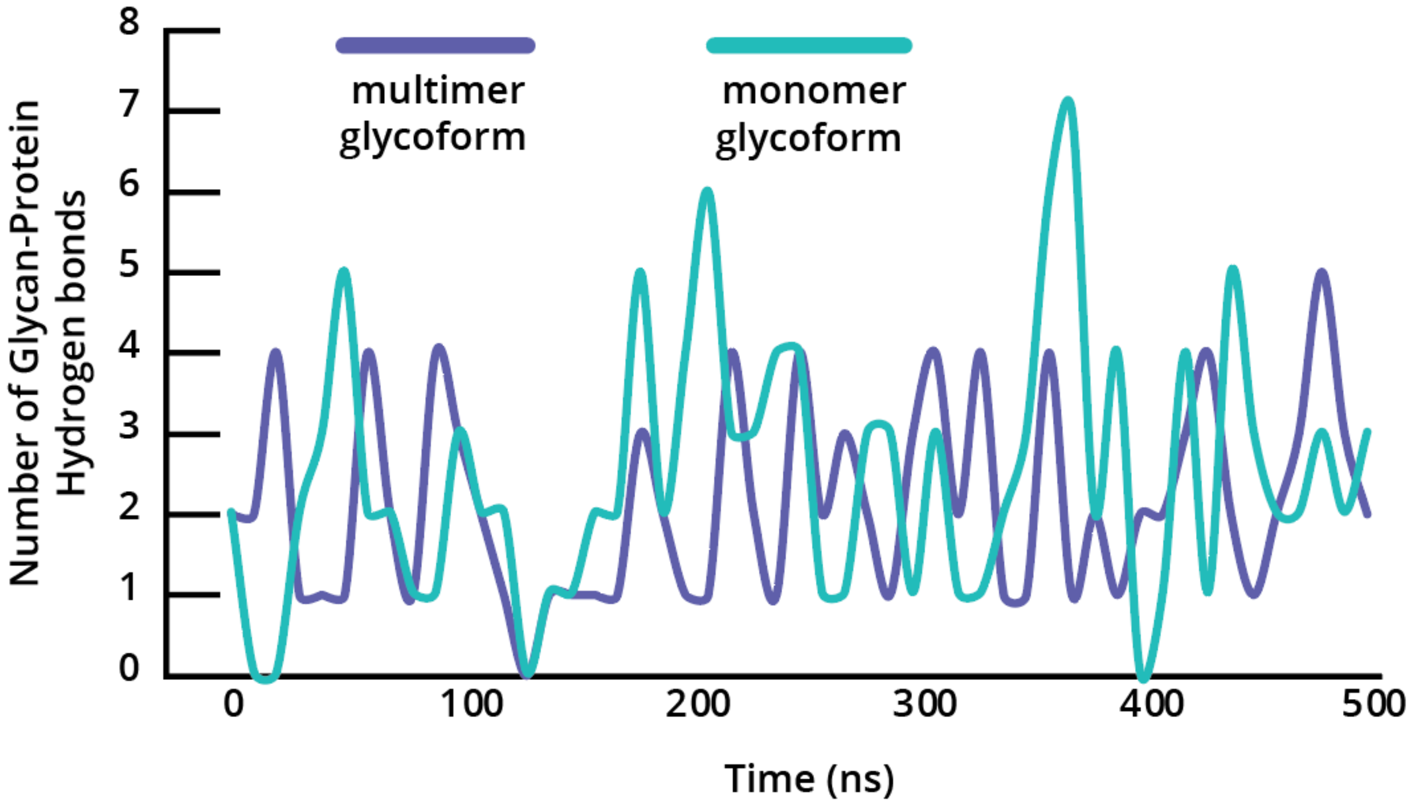


**Supplementary Figure 3. Intra-glycan-protein H-bonds over time**

A 500 ns production step was executed on GROMACs and the number of intra-glycan-protein H-bonds over time was extracted from the simulation. The number of H-bond interactions between glycans and the protein backbone are shown on the y-axis and the time is shown (in ns) on the x-axis for multimeric (blue) and monomeric (teal) lacritin.
